# Endurance exercise remodels pulmonary vein sleeve myocytes and promotes a proarrhythmic atrial substrate

**DOI:** 10.1101/2025.02.12.638004

**Authors:** Luca Soattin, Leila Topal, Roman Tikhomirov, Daniele Lagomarsino-Oneto, Sami Al-Othman, Sushant Saluja, Tibor Hornyik, Zoltán Husti, Jenő Antal Pintér, Aiman Saleh A Mohammed, Matthew Smith, Alice J Francis, Megan McKie, Eleonora Torre, Alexandra Polyák, Attila Farkas, Bo Hjorth Bentzen, Bernard Keavney, Norbert Nagy, Norbert Jost, Barbara Casadei, Matteo Elia Mangoni, Mark Richard Boyett, András Varró, Gwilym Matthew Morris, István Baczkó, Alicia D’Souza

**Affiliations:** Department of Biomedical Sciences, University of Copenhagen, Copenhagen, Denmark; Division of Cardiovascular Sciences, University of Manchester, Manchester, United Kingdom; Department of Pharmacology and Pharmacotherapy, University of Szeged, Szeged, Hungary; National Heart and Lung Institute, Imperial College London, London, United Kingdom; Department of Civil, Chemical and Environmental Engineering, University of Genoa, Genoa, Italy (now at Institute of Marine Sciences {ISMAR}, National Research Council {CNR}, Lerici, Italy); Doctoral School for Multidisciplinary Medicine, Albert Szent-Györgyi Medical School, University of Szeged, Szeged, Hungary; Institut de Génomique Fonctionnelle, Université de Montpellier, CNRS, INSERM, Montpellier, France; Department of Internal Medicine, Albert Szent-Györgyi Medical School, University of Szeged, Szeged, Hungary; Manchester University NHS Foundation Trust, Manchester Academic Health Science Centre, United Kingdom; NIHR Manchester Biomedical Research Centre, United Kingdom; HUN-REN-SZTE Research Group of Cardiovascular Pharmacology, Szeged, Hungary; Competence Centre of Pharmaceutical and Medical Device Developments of the Life Sciences Cluster, Centre of Excellence for Interdisciplinary Research, Development and Innovation of the University of Szeged, Hungary; Faculty of Life Sciences, University of Bradford, Bradford, United Kingdom; Department of Cardiology, John Hunter Hospital, Newcastle, Australia

## Abstract

**BACKGROUND:** Atrial fibrillation (AF) susceptibility is heightened in endurance athletes but the underlying mechanisms are incompletely understood. Because pulmonary vein (PV) myocyte triggers are critical determinants of AF, we investigated PV electrophysiological remodelling in animal models of the athlete’s heart.

**METHODS:** The following experiments were performed in canines and mice after 16 or 6 weeks, respectively, of daily exercise training (ExT), and compared to sedentary (Sed) controls: ECG recording, echocardiography, pharmacological autonomic block, extrastimulus pacing, multielectrode array mapping, monophasic and intracellular action potential (AP) recording with custom-designed pattern recognition analysis, histology, RNAseq and spatial *in situ* transcriptomics.

**RESULTS:** AF propensity was significantly increased in ExT animals. Mapping studies identified heightened rotational activity in the PV-left atrial (LA) junction of ExT *vs*. Sed canines *in vivo,* and enhanced automaticity, triggered activity and AP duration variability *ex vivo* in ExT canines and mice. Intracellular recordings in mouse PV cardiomyocytes determined at least six AP subtypes with increased frequency of pacemaker-like APs in ExT PV, concomitant with increased expression of pacemaking HCN4, Ca_v_1.3 and Ca_v_3.1 channels. PV spontaneous excitability was also significantly enhanced. Subcellular resolution spatial transcriptomics in mouse PV-LA identified diffuse ion channel remodelling and activation of established AF-promoting pro-inflammatory and pro-fibrotic cytokines and chemokines in ExT PV cardiomyocytes. Conduction slowing in the ExT PV-LA junction was attributable to: gap junction remodelling, reduced Na^+^ channel expression and increased extracellular matrix deposition with enhanced myofibroblast number and proximity to PV cardiomyocytes.

**CONCLUSIONS:** Endurance exercise elicits proarrhythmic electro-anatomical remodelling of the PV-LA junction with enhanced pacemaking ion channel expression and immune-inflammatory pathway activation in PV myocytes as prominent contributors.

**CLINICAL PERSPECTIVE:** *What is new?:*

- This work is the first demonstration that endurance training results in proarrhythmic electrophysiological remodelling of PV sleeve myocytes and extracellular matrix deposition in the PV-LA junction.
- We register electrical and molecular heterogeneity of the PV-LA junction at single cell and subcellular resolution, and for the first time identify the molecular events that underlie increased proarrhythmic activity of the trained PV. These include enhanced pacemaking ion channel expression (e.g., HCN4, Ca_v_1.3, and Ca_v_3.1), pro-inflammatory cytokine activation (e.g., TNFα, IL-6), increased myofibroblasts and extracellular matrix deposition.

*What are the clinical implications?:*

- We identify the molecular determinants of PV proarrhythmic activity in the trained heart and present new therapeutic targets for AF prevention in athletes.
- Our findings provide mechanistic rationale for the efficacy of pulmonary vein isolation for AF in athletes.

## INTRODUCTION

Endurance athletes are among the healthiest members of society yet paradoxically are prone to a range of cardiac arrhythmias^1^ including bradyarrhythmias, atrioventricular (AV) block, paroxysmal atrial fibrillation (AF) and ventricular arrhythmia,^2,3^ which are associated with significant morbidity and mortality. AF in particular has received attention in recent years as the best-evidenced example of an exercise-induced arrhythmia, commonly diagnosed in middle-aged (40-50 years^4–12^), otherwise healthy athletes engaging in intense endurance training for >10 years.^4,5,13–15^ Meta-analysis of observational studies indicates a 2.5-fold increased lifetime risk of AF compared with sedentary individuals despite a lower prevalence of conventional AF risk factors.^16^ The underlying mechanisms, assessed in rodent models, include heightened vagal tone and atrial substrate remodelling characterised by inflammation, fibrosis, *I*_K,ACh_ and *I*_Ca,L_ remodelling and ryanodine receptor phosphorylation.^17–20^ However, it is well recognized that AF is predominantly initiated by ectopic foci originating in the myocardial ‘sleeves’-thin layers of cardiomyocytes that extend from the left atrium (LA) and merge into the pulmonary veins (PVs).^21^ Mapping studies in humans and in isolated canine preparations have demonstrated that the PV–left atrial (LA) junction is a substrate for re-entry, and plays a critical role in initiation as well as maintenance of AF.^22,23,24^ Moreover, electrical isolation of the PVs has recently been shown to be efficacious in restoring and maintaining sinus rhythm in athletes,^25^ highlighting the importance of understanding training-induced changes to the PV-LA junction as a first step to understanding the most appropriate treatment and preventative strategies for AF in this subpopulation. Here we investigated PV myocytes sleeves and PV-LA junction remodelling in established canine and murine models of the athlete’s heart.

## METHODS

Full methodological details are provided in the Supplemental Material. Data supporting the findings of this study are available from the corresponding author upon reasonable request. Canine procedures were approved by the Ethical Committee for the Protection of Animals in Research at the University of Szeged, Hungary (approval numbers: I-74-15-2017, I-74-24-2017) and the Ministry of Agriculture and Rural Development (XIII/509/2022), in accordance with Directive 2010/63/EU. Exercise training in canines was performed as described previously.^26^ Murine studies were approved by the University of Manchester, complying with the UK Animals (Scientific Procedures) Act 1986. Briefly, ExT mice underwent 16 weeks of intensity-controlled interval training alternating between 8 min at 85-90% VO_2max_ and 2 min at 50-60% VO_2max_ for 60 min twice per day for 5 days per week. Echocardiography and ECG recordings were performed at baseline and post-training using previously established protocols.^26,29–32^ Canine open-chest multielectrode array (MEA) mapping, atrial monophasic action potential (MAP) recording and arrhythmia provocation studies were conducted under terminal anaesthesia. Murine Langendorff-perfused hearts underwent programmed electrical stimulation to assess atrial arrhythmia susceptibility. Canine and murine PV-LA preparations were isolated and superfused with oxygenated Tyrode’s solution for MEA and MAP recordings. Intracellular AP recordings were performed in the mouse common PV using sharp microelectrodes and analysed with custom-designed pattern recognition software incorporating machine learning. RNA sequencing (RNA-seq) and Xenium *in-situ* spatial transcriptomics were used to analyse differential gene expression, cell-type proportions, and spatial organization of the PV.

### Statistical analysis

*A priori* power calculations were performed to determine sample sizes (80% power, 95% confidence). Normally distributed data were analyzed using unpaired Student’s t-tests (with Welch’s correction for unequal variance), while non-parametric comparisons used the Mann–Whitney U test. χ^2^ tests assessed categorical variables such as arrhythmia inducibility. RNA-seq data were analyzed using DESeq2, and Xenium spatial transcriptomics employed Squidpy. Statistical significance was set at p<0.05.

## RESULTS

### Endurance training enhances AF vulnerability in canines

We previously reported on a 16-week high-intensity canine endurance training protocol that reproduced canonical structural and electrophysiological changes seen in human elite endurance athletes.^26^ Here, 7 canines that underwent this training protocol (ExT) and 7 sedentary (Sed) animals were studied. ECG recordings (**Figure 1A**) in conscious and unrestrained animals demonstrated a significant resting bradycardia following the training regimen, alongside prolongation of the PQ interval, QRS duration and the QT interval (**Figure 1B**). P wave duration was unchanged between groups (Sed, 44.92±3.9 ms; ExT, 55.04±2.4 ms, p=0.24). In line with our previous study,^26^ comparison of echocardiography parameters in ExT canines before (Week 0, pre) and after the training regimen (Week 16, post) confirmed increase in left ventricular posterior walls diameters (7.6±0.24 mm pre *vs*. 8.68±0.38 mm post, p=0.03, Paired t test) and increased left ventricular mass (64.6±4.3 gm pre *vs*. 77.2±3.8 gm post p=0.03, Paired t test) denoting training-induced left ventricular hypertrophy. Atrial dimensions did not vary significantly between groups. *In vivo* electrophysiological studies were performed to assess whether the training regimen impacted susceptibility to AF. **Figure 1C** shows representative surface MAP recordings from the left atrium during programmed electrical stimulation at 800 beats per minute (BPM) for 10 s. Comparison of the percentage of successful (AF-triggering) attempts out of 10 inductions demonstrated that training heightened vulnerability to AF (22.9±6.8 % *vs*. 57.1±12.3 %, **Figure 1C**, right panel). The mean duration of AF evoked by rapid atrial pacing in ExT dogs was 3.15±1.9 s, which was significantly longer than in the Sed group (1.03±0.5 s, **Figure 1D**, left panel). Similarly, rapid pacing in the presence of the cholinergic agonist carbachol (5 µg/kg) elicited AF with average AF duration 1.9±0.3 s in Sed versus 6.1±0.9 s in ExT animals (**Figure 1D, right panel**). It is concluded that in canines, as in humans, sustained high-intensity endurance exercise enhances AF vulnerability. Our model therefore constitutes a clinically relevant system for studying AF pathophysiology in endurance athletes.

**Figure 1.**
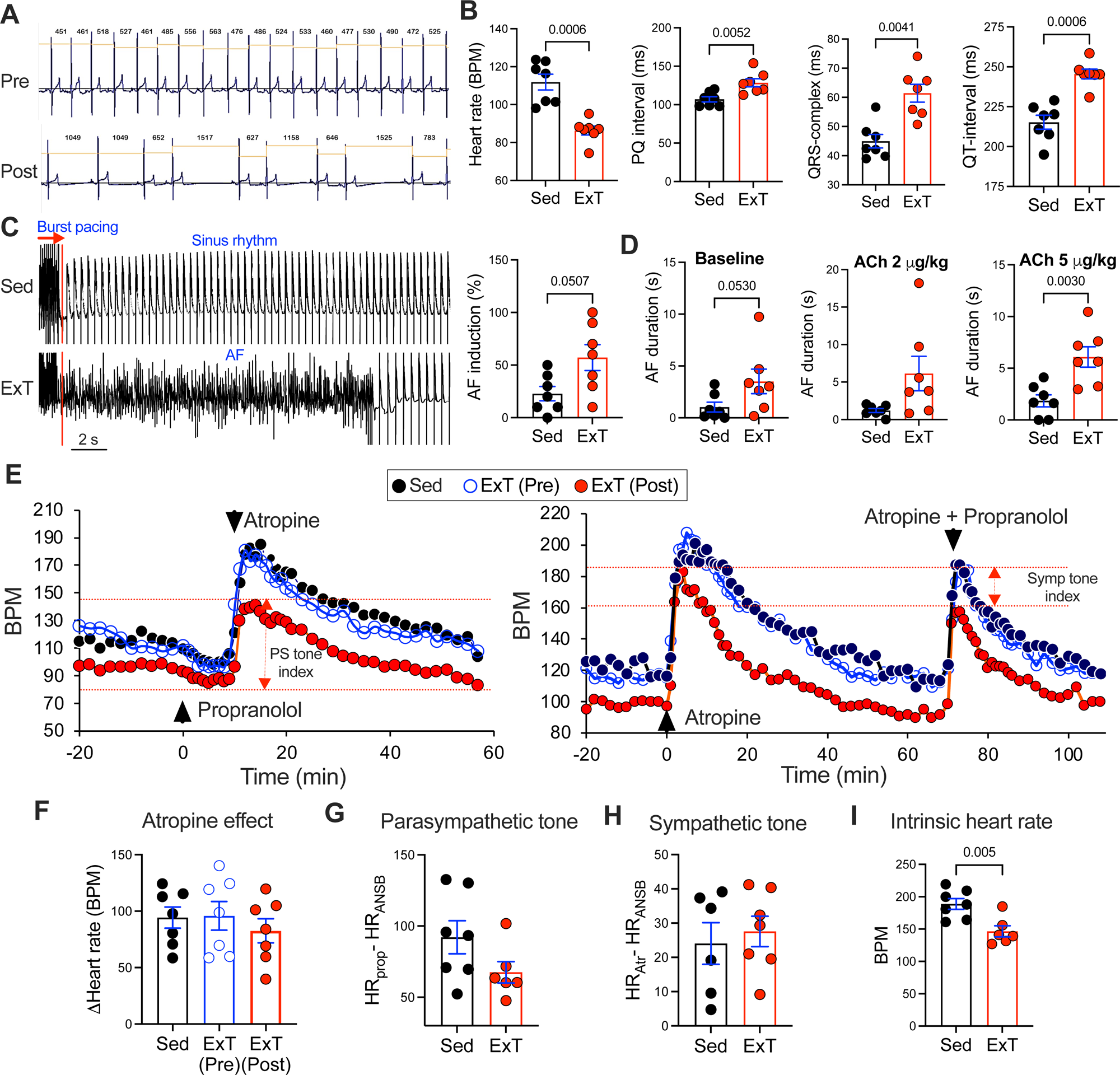
Athletes heart phenotype and enhanced AF susceptibility in ExT canines. **A**, Representative ECG recordings in an ExT dog prior to commencement of the training regimen (Pre) and on completion of training at week 16 (Post). **B**, ECG intervals in Sed (n=7) and ExT (n=7) canines. **C** (left), Representative monophasic action potential recordings during epicardial burst pacing (10 s, 800 beats/min, 3 x diastolic threshold) in Sed and Ext canines. Red line indicates cessation of burst pacing. **C** (right), AF inducibility, given as % successful AF induction attempts from a total of 10 trials of burst pacing in Sed (n=7) and ExT (n=7) canines. **D,** AF duration at baseline (left panel), in the presence of a bolus dose of acetylcholine delivered intravenously at 2 µg/kg (middle panel) and 5 µg/kg (right panel) in Sed (n=7) and ExT (n=7) canines. **E**, Mean heart rate recordings in conscious and unrestrained Sed (n=7), ExT (Pre, week 0, n=7) and ExT (Post, week 16, n=7 canines at baseline and during sequential sympathetic and parasympathetic block (left panel) and during sequential parasympathetic and double autonomic block (ANSB) (right panel). **F,** Maximal heart rate response to atropine in Sed (n=7), ExT (Pre, week 0, n=7) and ExT (Post, week 16, n=7) canines. **G-H,** Parasympathetic (G) and Sympathetic (H) tone in Sed (n=7) and ExT (n=7, week 16) canines. **I**, Heart rate under complete pharmacological block in Sed (n=7) and ExT (n=7, week 16) canines. Mean ± SEM given here and in all bar graphs. P values computed by Mann Whitney Rank sum test.

Because alterations in cardiac electrical function in athletes are commonly attributed to heightened parasympathetic tone, changes in autonomic tone were assessed by investigating the heart rate response to sequential pharmacological block of parasympathetic and sympathetic nervous input^17^ with atropine (0.04 mg/kg) and propranolol (0.2 mg/kg) respectively. Doses of autonomic blockers were selected based on previous canine studies^27,28^ and verified in prior pilot experiments with agonist challenge. **Figure 1E** gives mean heart rate measurements over the recording period in conscious and unrestrained Sed and ExT dogs, demonstrating that the heart rate response to atropine did not differ between Sed and ExT groups or on comparison of pre- and post-training changes in the ExT group (**Figure 1F**). As such, changes to parasympathetic tone in ExT animals were not detectable in this study (**Figure 1G)**. Sympathetic tone was also unaltered between groups (**Figure 1H**) but conversely, intrinsic heart rate, i.e. heart rate in presence of both atropine and propranolol, was significantly reduced by exercise (188.9 ± 8.3 BPM for Sed *vs*. 146.6 ± 8.6 BPM for ExT dogs at 16 weeks, **Figure 1I**). The spontaneous beating rate of isolated right atrial preparations was also reduced following the training regimen (**Supplemental Figure S1**), in line with training-induced intrinsic remodelling of the sinus node independent of vagal tone that we have previously described in human athletes^29^ and in animal models.^26,30–32^

### Endurance training modifies PV-LA conduction patterns and PV excitability

High density MEA mapping of the left inferior PV and PV-LA junction with concomitant MAP recording of the left inferior PV and left atrial appendage (LAA) were performed to investigate PV activation and conduction patterns *in vivo* (**Figure 2A**). During sinus rhythm, spontaneous ectopic beats that originated in the PV and propagated to the LA were observed sporadically in ExT but not Sed hearts (**Figure 2B**). Representative data from MEA mapping in ExT hearts given in **Figure 2B** shows that such ectopic beats abruptly change conduction trajectories and propagate from the PV to the LA. Under programmed electrical stimulation from the LAA (150 BPM) distinct lines of block were seen in the LA-PV junction in all ExT (**Figure 2C**) but not Sed hearts, corresponding to a significant increase in local activation time and decrease in conduction velocity in ExT hearts **(Figure 2D-E).** Interestingly, significant changes to radial propagation patterns of conducted stimuli were also discernible between ExT (162 ± 24°) compared to Sed (96 ± 14°) groups **(Figure 2F)** suggesting that exercise training alters activation patterns within the PV-LA junction. PV action potential (AP) parameters including AP duration at 90% repolarisation (APD_90_) did not differ between groups (130.7 ± 18.4 ms in Sed *vs*. 143.2 ± 21.1 ms in ExT; p>0.05) Compared to Sed PV, a trend towards increased short-term APD variability, commonly associated with a proarrhythmic substrate, was observed in ExT PV but the difference did not reach statistical significance (**Supplemental Figure S2**).

**Figure 2.**
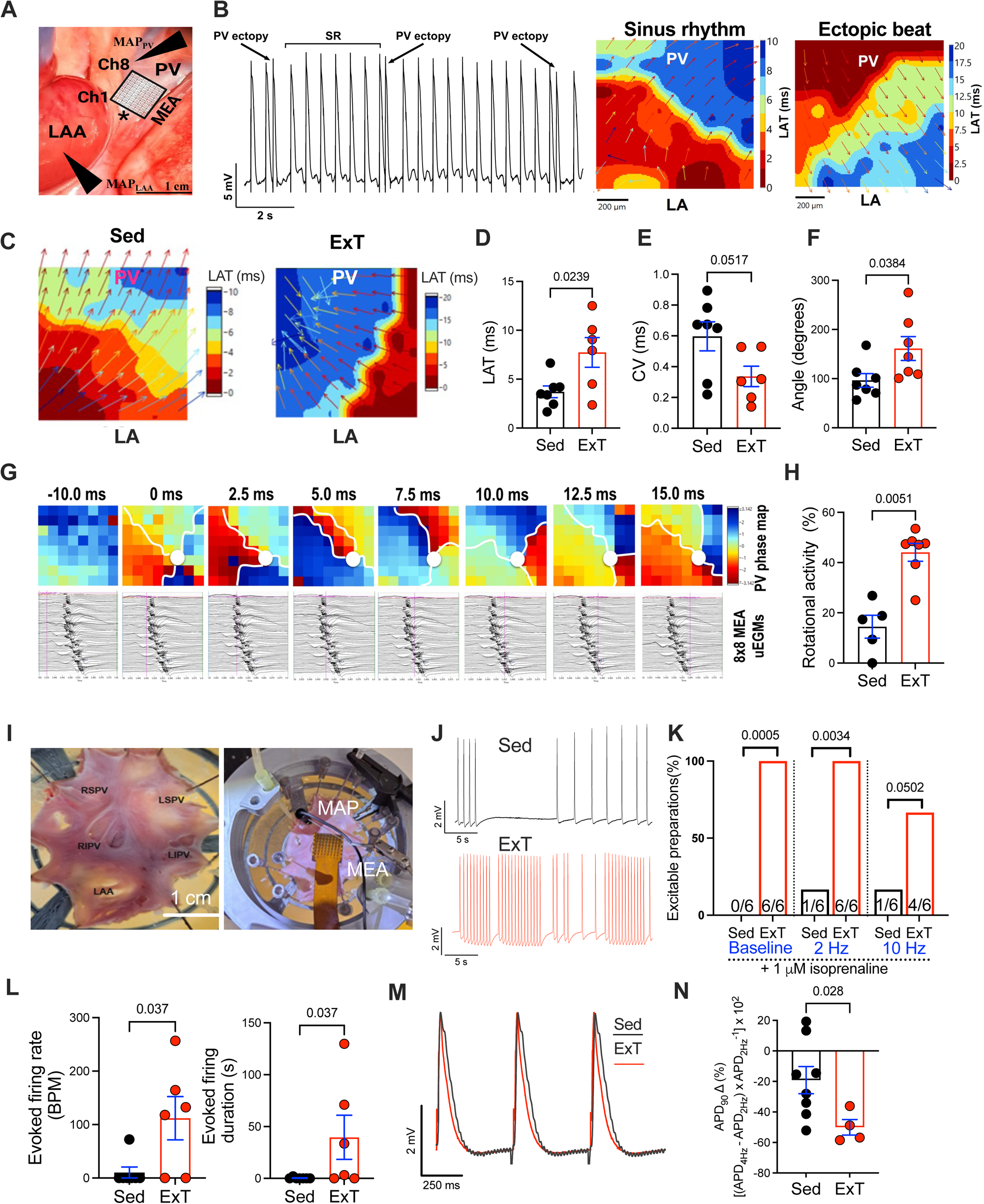
Exercise training results in altered conduction patterns, increased rotational activity and intrinsic excitability in the PV-LA junction in canines. **A**, Placement of the 64Ch-MEA on the epicardial surface of the PV-LA junction (represented by asterisk) during *in vivo* open chest experiments in canines. Ch1 and Ch8 indicate the orientation of the MEA. MAP electrodes were positioned on the LAA or on the PV (arrows). **B**, Representative concomitant MAP (left panel) and MEA recordings (right panel) of spontaneous ectopic beats in an ExT PV propagating to the LA. Isochronal activation maps denote local activation time, in which earliest activation is in red and latest is in blue. Arrows represent local direction of propagation. **C,** Representative PV**-**LA isochronal maps during 2.5 Hz pacing from the LA appendage in Sed and ExT canines. Note isochronal crowding and prolonged local activation times in ExT PV-LA. **D-F,** Summary data from experiments in C denoting local activation time (D), conduction velocity (E) and direction of radial propagation (F) in Sed (n=7) and ExT (n=7) canines. **G**, Representative recording of rotational activity in the canine PV-LA junction. Top panel, phase maps at baseline (-10 ms), and during a complete rotational re-entrant circuit during an AF episode. Circle indicates the rotor core, also defined by the LAT map showing a region of isochronal crowding and wavefront curvature, leading to slowing in conduction (>0.03 m/s). Red indicates the depolarisation phase of the wavefront, and blue, repolarisation phase. The wavefront in the PV propagates around the indicated centre, generating rotational activity. Bottom panel, Pink cursor marks depolarisation and repolarisation events in channels in the time window specified. Images at 2.5 ms intervals shown. **H**, Rotational activity occurence (%) relative to total number of beats analysed during 5 s of AF evoked by burst pacing in the presence of 5 µg/kg acetylcholine in Sed (n=5) and ExT (n=7) canines. **I**, Canine PV-LA preparations and position of MAP and MEA electrodes in the perfusion chamber. **J**, Representative MAP recordings of spontaneous PV ectopic activity in isolated PV-LA preparations from Sed and ExT canines following cessation of pacing (2 Hz). **K**, Summary data showing percentage of PV-LA preparations in which PV ectopic activity could be evoked using 1 µM isoproterenol or on electrical stimulation for 3 s at 2 Hz and10 Hz in Sed and ExT canines. Number of preparations analysed per treatment are given in bar charts. **L**, Rate (left panel) and duration (right panel) of post pacing evoked firing in Sed (n=7) and ExT (n=6) canines following 10 Hz burst pacing in the presence of 1 µM isoproterenol. **M,** Representative MAP recordings in Sed and ExT PV indicating shorter APD90 duration in ExT preparations. **N,** APD90 shortening relative to pacing rate at 2 Hz and 4 Hz in the presence of 1 µM isoproterenol in Sed (n=8) and ExT (n=4) PV. P values computed by Mann Whitney rank sum test (panels **D-F**, **H**, **L** and **N**) or χ^2^ test (panel **K**).

During triggered AF, distinct spatiotemporal activation and conduction patterns were observed between groups. Rotational activity, defined as sequential clockwise or counter-clockwise activation contours (isochrones) around a centre of rotation emanating outward, was apparent in MEA recordings in 7/7 ExT and 5/7 Sed hearts (**Figure 2G,H**). Phase analysis of AF events evoked by burst pacing in the presence of 5 µg/kg carbachol demonstrated that rotor activity (calculated as rotors/total number of beats during 5 s of AF for each animal) was significantly more frequent in ExT PVs compared to Sed PVs (**Figure 2H**, Sed 14.5 ± 4.6%; ExT 44.1 ± 3.5%, p<0.01). An example of rotational activity in an ExT PV during AF based on local activation time phase maps covering one cycle is illustrated in **Figure 2G**. In both groups, rotational activity was typically sustained for at least one full rotation in the anti-clockwise direction before breaking into less organized wavelets. PV focal activation, defined as centrifugal isochrones from an origin in the PV, was frequently observed in the ExT group whereas in the Sed group the PV was typically passively activated by a wavefront arising in the LA. To investigate intrinsic changes to PV excitability and conduction, denervated PV-LAA preparations were rapidly dissected on termination of *in vivo* experiments and then superfused, and MAP and MEA recordings performed following a one-hour stabilisation period (**Figure 2I)**. 6 out of 7 preparations in both groups were viable and investigated further. All preparations were quiescent at baseline, but interestingly β-adrenergic stimulation with 1 µM isoproterenol triggered APs in ExT preparations only (**Figure 2K**, 0/6 Sed preparations *vs*. 6/6 ExT preparations). When paced at 2 Hz (3 s) from the right superior PV, a higher proportion of ExT PVs generated post pacing spontaneous activity (100%, 6/6 preparations) compared to Sed PVs (16.7%, 1/6 preparations, **Figure 2J-K**) in the presence of 1 µM isoproterenol. Similarly rapid pacing (10 Hz, 3 s) in the presence of 1 µM isoproterenol stimulated spontaneous activity in more ExT PVs (66.6%, 4/6 preparations) than Sed PVs (16.7%; 1/6 preparations) on pacing cessation (**Figure 2K**). Moreover, spontaneous firing bursts on cessation of pacing in ExT PV were significantly longer and of significantly higher frequency than in Sed PV preparations (**Figure 2L**). Similar to *in vivo* MAP recordings, AP duration at 90% repolarisation (APD_90_) at a pacing frequency of 2 Hz demonstrated a trend towards reduction in the trained group at baseline (**Figure 2M**, 176 ± 18 ms in Sed *vs*. 157 ± 7 ms in ExT; P>0.05, Mann Whitney rank sum test) and in the presence of 1 µM isoproterenol (132 ± 13 ms *vs*. 116 ± 8 ms; P>0.05, Mann-Whitney rank sum test). Furthermore, the rate dependence of APD was greater after exercise training (**Figure 2N**). Taken together, these data provide the first direct evidence that sustained endurance training modifies activation patterns and rotational activity in the LA-PV junction, and also enhances triggered activity and intrinsic excitability of PV sleeve myocytes.

### PV electrophysiological remodelling in a mouse model of training-induced AF

The findings above prompted a detailed exploration of endurance training-induced PV-LA remodelling. Male C57BL6/J mice were subjected to 6 weeks of treadmill exercise (ExT), running for 2 h/day, 5 days/week with intervals of 8 min at 85-90% VO_2max_ and 2 mins at 50-60% VO_2max_. This protocol has previously been shown to recapitulate human cardiac and skeletal muscle adaptation to exercise^33^; full details in **Supplemental Methods**. Mice in the sedentary (Sed) group were acclimatised to the treadmill apparatus by placing on a flat treadmill at 0.15 m/s for 15 min daily. **Figure 3A** shows that at the end of the 6-week training period VO_2_ scaled for body mass (reported as ml/min/kg^-0.75^) increased by 20.5% in ExT *vs*. Sed mice. Echocardiographic measurements revealed a 41.3% increase in left ventricular relative wall thickness (**Figure 3B**, right panel), and significant increases in diastolic left ventricular internal diameter and posterior wall diameter (**Supplemental Table S1**). Similarly left atrial anteroposterior diameter was also enhanced by training (**Figure 3C**). ExT mice presented with further canonical features of the athlete’s heart including sinus bradycardia and prolongation of the PR interval (**Figure 3D**). These changes persisted on complete pharmacological autonomic blockade with 0.5 mg/kg atropine and 1 mg/kg propranolol, indicating intrinsic electrophysiological remodeling as seen in ExT canines (**Figure 1I** and **Supplemental Figure S1**) and as previously described by our group in other models of endurance training.^29–32^ AF inducibility was assessed in the isolated Langendorff-perfused heart using an extrastimulus pacing protocol^34^ in the presence of the cholinergic agonist carbachol (1 µM). As anticipated, propensity to pacing-induced atrial tachyarrhythmia, defined as rapid and chaotic electrical activity exceeding 10 s,^19^ was significantly increased in ExT mice (**Figure 3E**, 0% in Sed *vs*. 57.14% in ExT). In sum, this mouse model mimicked human responses to sustained endurance exercise, and was deemed an appropriate platform to explore training-induced structural, electrophysiological and molecular remodelling of the PV-LA junction.

**Figure 3.**
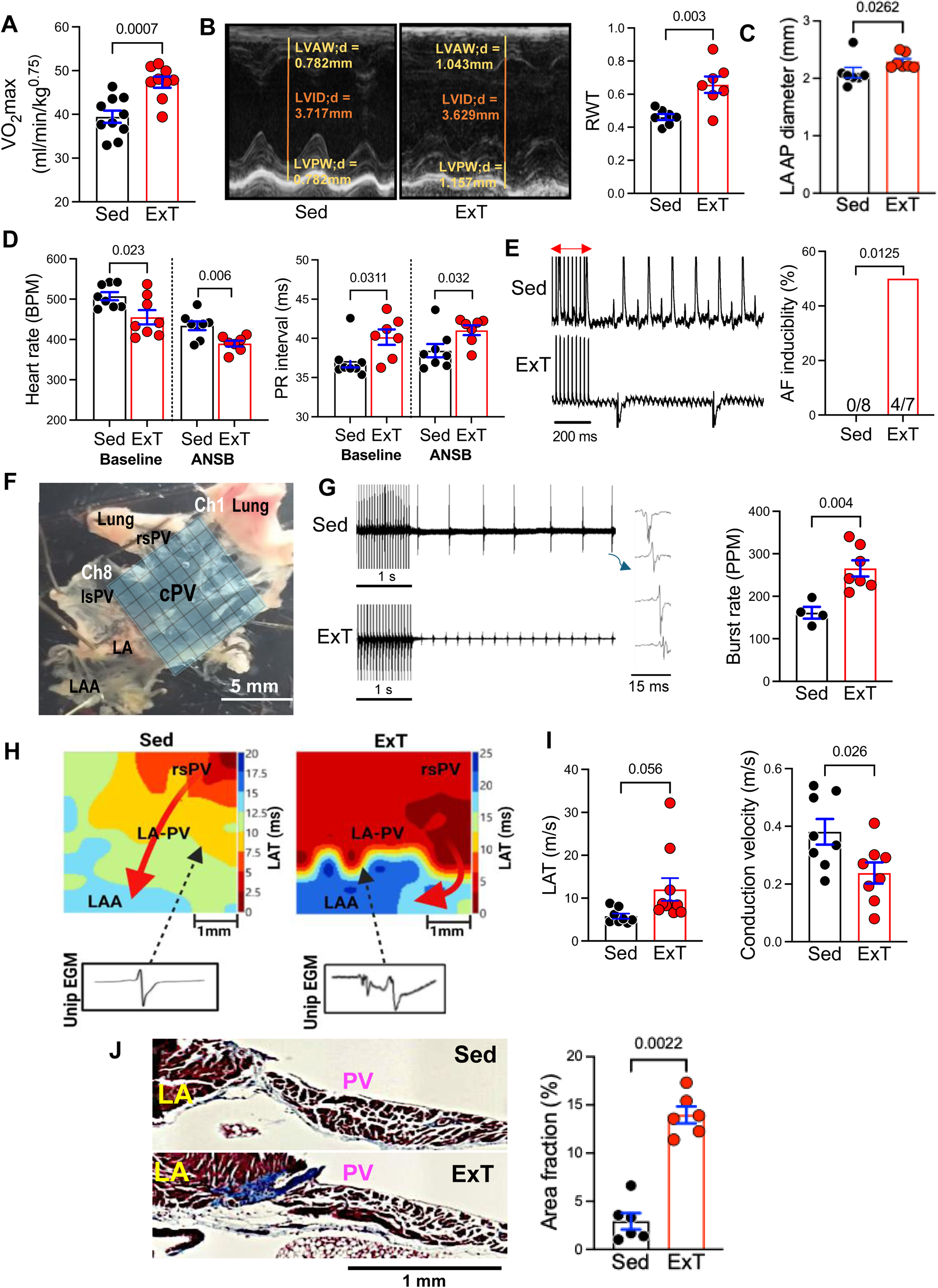
Athletes heart phenotype, enhanced AF susceptibility and PV-LA junction remodelling in ExT mice. **A**, Normalised VO_2max_ of Sed (n=8) and ExT (n=8) mice following 6 weeks of intensity-controlled interval treadmill training. ExT mice ran 60 min per day, 5 days per week for 6 weeks in intervals of 8 min at 60-80% VO_2max_ and 2 min at 40-50% VO_2max_. **B** (left), Representative M-mode echocardiogram obtained from short axis parasternal view of the left ventricle (LV) in Sed (n=8) and ExT (n=7) mice. **B** (right), LV relative wall thickness (RWT) in Sed (n=8) and ExT (n=7). **C**, Left atrial anteroposterior (LA AP) diameter in Sed (n=8) and ExT (n=7). **D**, Heart rate (left) and PR interval (right) obtained during anaesthetised 3-lead ECG recordings in Sed (n=8) and ExT (n=7) mice at baseline and during pharmacological autonomic block (ANSB) with atropine (0.5 mg/kg) and propranolol (1 mg/kg). **E** (left), Representative ECG recordings during extra stimulus pacing in Langendorff-perfused Sed and ExT hearts in the presence of 1 µM carbachol. Red arrow denotes pacing train. **E** (right), Summary data from experiments in E, (left) quantifying % susceptibility to pacing-induced atrial tachyarrhythmia in Sed and ExT groups. N numbers per group shown in bars. **F**, Mouse PV-LA preparations with schematic representation of MEA position. Channel numbers (Ch) indicated to aid orientation. rsPV, right superior PV; lsPV, left superior PV; cPV, common PV. **G** (left), Representative traces during 10 Hz pacing (3 s) from the rsPV and ensuing spontaneous unipolar electrograms (uEGMs) in Sed and ExT preparations. Inset shows individual EGMs **G,** (right) Summary data from experiments in left panel showing frequency of evoked EGMs in Sed (n=4) and ExT (n=7) PV. PPM, pulses per min. **H**, Representative MEA isochronal maps of activation time in Sed and ExT preparations. Representative uEGMs at selected locations given in inset. Typical biphasic uEGMs observed at LA-PV junction in Sed PV while in ExT PV uEGMs were often fractionated or displayed double potentials. Arrows indicate wavefront propagation path. **I**, Mean local activation time (left) and mean conduction velocity in Sed (n=8) and ExT (n=10) preparations calculated from experiments given in panel H. **J** (left), Representative Masson’s trichrome staining of 10 µM sections of the PV-LA junction in Sed and ExT preparations**. J** (right), Summary data of ECM area fraction in Sed (n=6) and ExT (n=7) preparations computed from experiments in left panel. Four slides from each preparation were analysed. P values computed by Student’s t test (panels **A-D, G, I**) Mann-Whitney rank sum test (**J**), or χ^2^ test (**E**).

Electrophysiological study of the isolated and denervated PV-LA preparation was first performed using high-density MEA mapping of the endocardial surface (**Figure 3F**). Programmed electrical stimulation (1 s, 20 Hz, 2 x diastolic threshold) at the right superior PV ostium triggered a higher spontaneous pulse frequency on pacing cessation in ExT *vs*. Sed PVs indicating increased excitability of ExT PVs (**Figure 3G**), as observed in canines (**Figure 2K**). Analysis of isochronal activation maps from MEA recordings identified anisotropy in conduction from the site of earliest activation (SEA) in the PV-LA preparation, underscored by beat-to-beat variability in preferential conduction directions within the same burst (**Supplemental Figure S3A**). Subtle changes to location of the SEA were appreciable between groups at baseline and in the presence of 10 µM noradrenaline (See SEA profile in **Supplemental Figure S3B-C).** For example, 10 µM noradrenaline resulted in an inferior shift of the SEA from the rsPV to the LA-PV junction in 50% ExT preparations but only 25% of Sed preparations. Considering conduction times, ExT-PV preparations showed a significant increase in local activation times relative to Sed-PVs (**Figure 3H-I**). ExT preparations presented with an increased proportion of fractionated electrograms (indicative of conduction inhomogeneity) which corresponded to distinct lines of conduction block (**Figure 3H**) in the PV-LA junction (CV<0.1 m/s). As a consequence, conduction velocity was significantly prolonged in ExT *vs*. Sed PVs (**Figure 3I**), constituting a substrate conducive to re-entry. Extracellular matrix (ECM) remodelling is known to promote heterogeneity of conduction and pathways facilitating micro-reentry and macro-reentry^35^ and thus, Masson’s trichrome staining on 10 µM thick sections of the PV-LA preparation was performed to assess whether slowed conduction in ExT PVs was attributable to enhanced ECM deposition. **Figure 3J** shows that ECM area fraction was significantly increased in ExT PV, ranging from 2.9 ± 0.8% in Sed to 11.4 ± 0.8% in ExT preparations. ECM deposition in ExT PV was particularly pronounced at the interface between the PV and the LA, concomitant with conduction slowing in this region (**Figure 3H**).

Next, to characterise PV electrophysiological remodelling at the cellular level, intracellular APs were recorded using high resistance sharp microelectrodes (3M KCl, 20-50 MΩ) in endocardial PV regions of intact PV-LA preparations from ExT (n=9) and Sed (n=12) mice. In these experiments robust spontaneous activity with typical^36^ burst firing patterns and a range of APs including atrial-like APs, APs with early afterdepolarisations, and ‘pacemaker’ like APs with diastolic depolarisation were observed in both groups (**Figure 4A**, see also **Figure S3A**). In contrast to previous work^37^ specific AP morphologies were not restricted to specific regions within the PV but instead appeared randomly distributed within the sleeve. It was thus apparent that PV myocytes are heterogenous populations with distinct kinetics, that also exhibited chaotic alterations between active (firing) and rest (non-firing) phases (**Figure 4B**). These characteristics posed significant challenges for quantification and comparison of ExT-induced changes to PV activity. To circumvent this issue, we developed a novel pattern recognition analysis pipeline integrating machine learning (**Figure 4C**) to automatically detect and quantify group differences in AP morphology, firing patterns and intra-burst frequency. A full description of the algorithm is given in **Supplemental Methods**. Briefly, we analysed ∼30 h of membrane potential recordings from Sed and ExT preparations. After pre-processing and evaluation of resting membrane potentials, we defined ‘active phases’ (segments of the signal in which the recorded voltage significantly exceeded the threshold for AP firing) permitting identification of AP spikes, and aggregation of spikes into bursts. Recordings from each preparation were split in subsets at three nested levels: chunks (defined as valid time-series of contiguous times), phases (defined as either bursts or rests) and single events within a particular active phase. At the single-event level we further defined the following variables: (**i**) shape class - stratified according to AP morphology using K-means clustering,^38,39^ (**ii**) APD (**iii**) AP amplitude and (**iv**) resting membrane potential. At the phase level, duration of bursts and rests and the number of events within a burst. These analyses resolved significant heterogeneity in AP properties and alterations in firing patterns in ExT *vs*. Sed PV myocytes and several interesting findings emerged: first, recorded APs were clustered into 6 different shapes (numbered 0-5; see **Methods** for further details), including atrial-like APs (shape class 0,1 and 5), pacemaker-like APs (APs with phase 4 depolarization, shape class 2) and APs with early afterdepolarisations (shape classes 3 and 4) (**Figure 4D**). Second, AP parameters broadly aligned with previous intracellular recordings in the mouse PV^36^ but within a given shape class, AP diastolic potential, amplitude and duration showed significant heterogeneity and non-gaussian distributions in most cases (APD variation given as an example in **Supplemental Figure S4**; see also **Supplemental Table S3** and **Supplemental Discussion**). Comparison of modal values (values occurring most frequently in the dataset) indicated shorter APDs in Type 0 atrial-like APs (**Supplemental Figure S4** and **Supplemental Table S3;** See also **Figure 2M)** and such AP heterogeneity is expected to favour occurrence of re-entry. Third, frequency of a given AP shape, defined as number of single events of a particular shape class revealed a distinct distribution of AP subtypes in ExT and Sed groups (**Figure 4E**) when analysed using a χ^2^ test with a stringent false discovery rate (FDR) correction <1x10^-10^. Importantly, the percentage of pacemaker-like APs increased from 6.69% in Sed PV to 12.53% in ExT PV (χ^2^ = 129.58, FDR = 6.18 x 10^-30^) likely explaining enhanced excitability in ExT preparations reported in **Figure 3G**. Atrial-like APs (shapes 0 and 1) varied in distribution between groups in which the frequency of shape 0 was increased by 17% (χ^2^ = 437.4, FDR = 1.4 x 10^-96^) and shape 1 was reduced by 26.69% in ExT PV (χ^2^ = 902.24, FDR = 2.2 x 10^-197^). Fourth, considering burst firing patterns, there was a prolongation in the duration of spontaneous bursts in ExT PV (0.9 ± 0.06 s in Sed *vs*. 1.4 ± 0.14 s in ExT, **Figure 4F** left panel) coupled with an increase in the number of APs per burst (**Figure 4F**, right panel). Burst patterns are considered in further detail in **Supplemental Figure S5**. Intermittence, defined as burst duration (*T*_active_) normalised to total phase duration (*T*_active_ + *T*_rest_) (**Figure 4G**) demonstrated markedly enhanced spontaneous ectopic activity in ExT animals, in that the time spent in active phases was increased in ExT *vs*. Sed preparations (Sed 0.36±0.08 vs. ExT 0.52±0.08). In ExT mice, the proportion of bursts with durations longer than 1.01 s and 4.01 s were greater than in Sed mice (**Supplemental Figure S5**). Taken together, these novel approaches demonstrated that sustained endurance exercise enhances pro-arrhythmic properties of PV myocytes, characterised by increased spontaneous activity, enhanced automaticity, APD changes in atrial-like APs and faster burst firing patterns, that when coupled to conduction slowing (**Figure 3I**) and ECM deposition in the PV-LA junction (**Figure 3J**) could predispose to the initiation and maintenance of AF.

**Figure 4.**
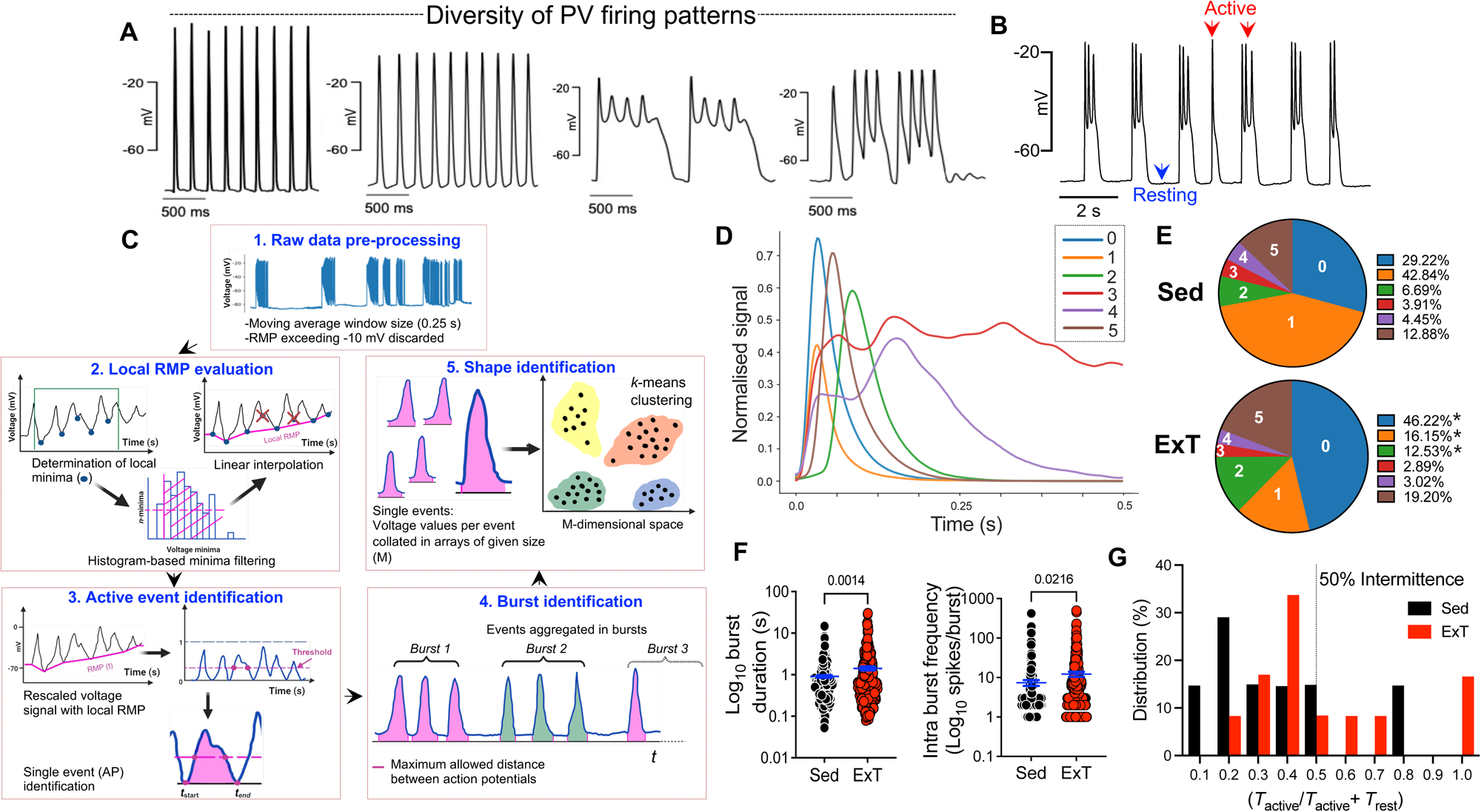
Dynamic pattern analysis defines PV AP heterogeneity and its modulation in the ExT heart. **A**, Representative intracellular APs recorded using sharp microelectrodes in mouse Sed PV preparations demonstrating exemplar AP subtypes in the PV. **B**, Representative intracellular APs in mouse Sed PV demonstrating typical intermittent burst firing patterns. **C**, Custom designed workflow for automated dynamic analysis of intracellular AP recordings from n=11 Sed and n=12 ExT PV: 1, Raw AP data were processed using a moving average window size of 0.25 s. In case of RMP drift (depolarisation exceeding -10 mV) fragments were manually disregarded. Resultant “temporal slices” were analysed automatically. 2, Local minima were identified by a moving window, a histogram system used to remove minima that did not represent take-off potentials, and linear interpolation used to find regular minima. 3, Interpolated voltage (V) minima were rescaled according to the local RMP, using a V/|RMP| ratio, enabling single event identification in the temporal slice. 4, Definition of bursts by aggregation of events. 5, AP shape determined by collecting voltage values for single events in arrays of given size M (e.g., V1, V2,…VM). Events presenting > M-values were truncated at the first M-value, while shorter events padded with their minimum recorded value. Each array was shifted and scaled to avoid separating events with similar shape but different RMP. K-means clustering grouped events with similar features. **D**, PV action potentials stratified into 6 subtypes (averaged shape) from analysis described in panel C. **E**, Pie charts demonstrating proportion of 6 detected AP subtypes computed as the fraction of events of a given shape in Sed (top panel) and ExT (bottom panel) preparations. **F**, Automated analysis of burst duration (left) and intraburst frequency (right) during active phases in Sed and ExT PV myocytes. **G,** Histogram showing distribution of intermittence in Sed and ExT PV. Values >0.5 indicate sustained burst activity. Value distribution as median, 25% and 75% percentile: Sed (0.27, 0.20, 0.46); ExT (0.43, 0.35, 0.67). Points denote discrete phases. P values derived from a Mann-Whitney rank sum test (**F,** left) or Student’s t test with Welch’s correction for unequal variance (**F,** right).

### Molecular signature of training-induced PV remodelling

For insight into the molecular signature that underpins ExT-induced PV remodelling, bulk RNAseq was performed in biopsies of the mouse common PV. **Figure 5A** shows that of the 55,401 transcripts measured, 3,590 transcripts were differentially expressed in ExT (n=7) *vs*. Sed (n=8) PVs of which 3,415 were upregulated and 2,382 were downregulated in ExT PV (p<0.05; FDR p<0.1). Among significantly upregulated transcripts was the ion channel subunits *Hcn4* (cardiac hyperpolarization-activated cation channel 4 underpinning *I*_f_ - a major determinant of diastolic depolarization in pacemaker cardiomyocytes) (**Figure 5B**). A subset of PV cardiomyocytes have been shown to express HCN channels^40^ and exhibit *I*_f_^41^ and this finding could explain the increased occurrence of pacemaker-like APs observed in ExT PV (**Figure 4D**). Furthermore, gene ontology analysis of differentially expressed genes revealed striking immune-inflammatory activation in ExT PV characterised by increased gene expression of several well-established cardiac pro-fibrotic and pro-inflammatory signalling mediators (**Figure 5B**) including TGFβ, NFkB, interleukin-6 (Il6), interleukin-11 (Il11), chemokine (C-X-C motif) ligands CXCLs 1,2,3,5 and 11 as well as cytokine receptors (e.g. TNFR1). This is important from a mechanistic perspective as there is a robust association between proinflammatory cytokine levels and AF progression in patients,^42^ proinflammatory cytokines Il6 and TNFα are chronically elevated in human athletes^43,44^ and specific contributions of cytokines/chemokines in atrial remodelling and AF have been determined.^19,45,46^ We hypothesised that ion channel changes and pro-inflammatory signalling described above were a consequence of altered cell type composition and expansion of macrophages that has been previously described in the atria of exercise-trained trained mice.^19^. To test this hypothesis, we deconvoluted and determined the cell type composition of each of the 16 bulk RNA-seq samples, leveraging published mouse PV myocyte single cell RNA-seq (scRNA-seq) data for estimation of cell type proportions.^47^ The expression levels of marker genes for 18 cardiac cell types (including 3 cardiomyocyte subtypes, 2 fibroblast subtypes, lymphoid and myeloid cells – see **Supplemental Methods** for full details) was input into CIBERSORTx^48^, a widely used cell-type deconvolution tool based on support vector regression to identify subpopulations and heterogeneity within a mixed cell population. While this approach did not reveal any statistically significant changes in immune cell types between Sed and ExT groups (**Supplementary Table S3**), significant changes in fibroblast (FB) subtypes FB1 and FB2, and a trend towards increase in the proportion of a cardiomyocyte subtype (CM2) was observed in ExT PV (**Figure 5C**). Because such alterations could mechanistically contribute to increased ECM deposition and altered AP subtypes (respectively) observed in ExT PV (**Figure 3J**, **Figure 4E**), we investigated these ExT-induced changes at single-cell resolution and in a spatially resolved manner using a state-of-the-art targeted *in situ* transcriptomics approach.

**Figure 5.**
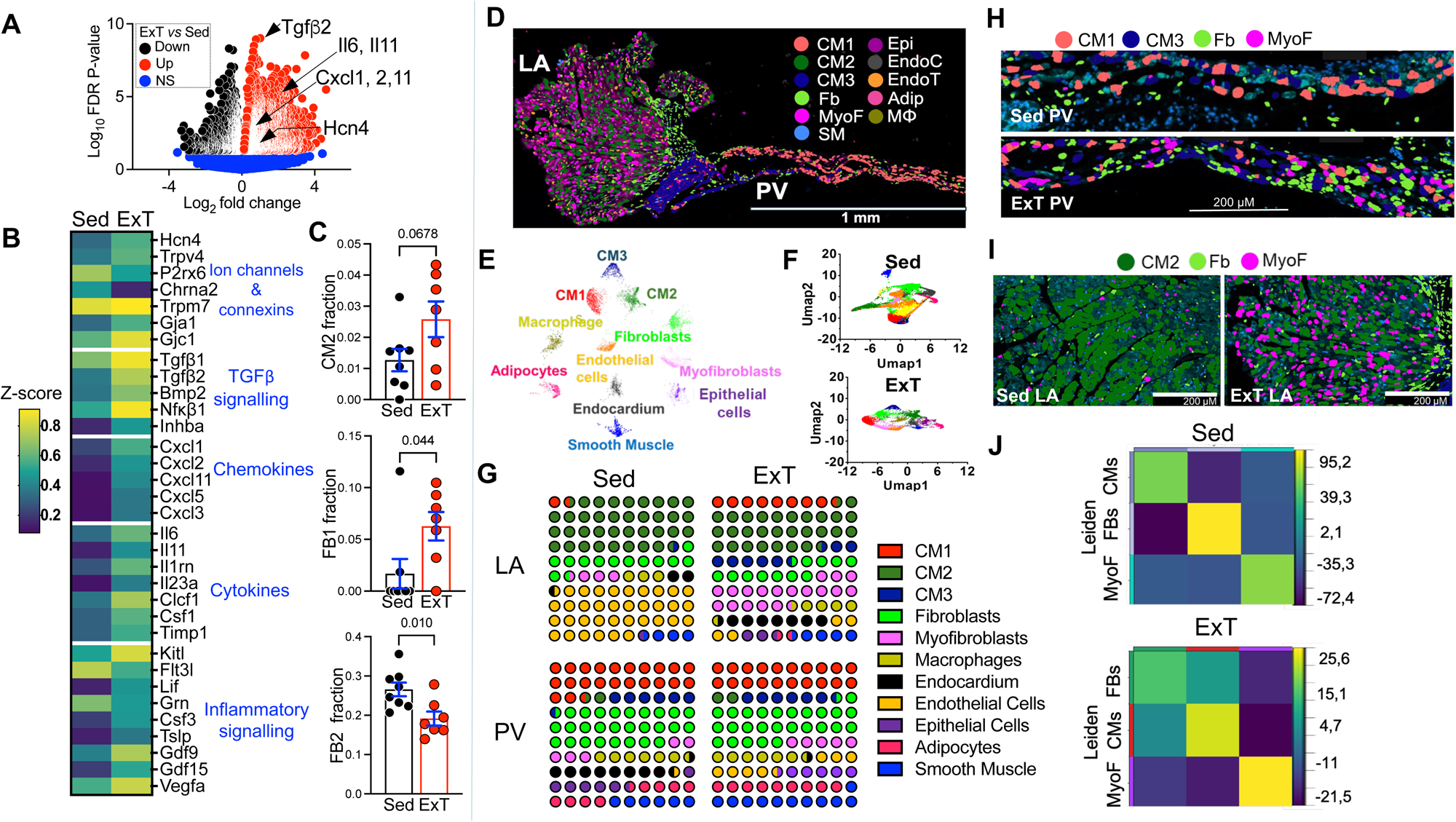
Molecular landscape of the PV-LA junction at subcellular spatial resolution. Volcano plots representing differentially expressed genes (DEGs) determined by RNAseq in 1 mm common PV biopsies of Sed (n=8) and ExT (n=7) mice. Red circles, significantly up-regulated genes (N = 3415, FDR < 0.05); black circles, significantly down-regulated genes (N = 2382, FDR < 0.05); blue circles – unaltered genes. Transcripts associated with inflammation (*Tgfβ2*, *Il6*, *Il11*, *Cxcl1*,*2*,*11*) and *Hcn4* transcript are highlighted. **B**, Heatmap displaying normalised mean values per condition (Sed or ExT) for significantly modulated genes (FDR < 0.05) grouped and annotated by their biological functions: Ion channels and connexins; TGFβ signaling; Chemokines; Cytokines and Inflammatory signalling. Data were z-scored by gene. **C**, Bar graphs demonstrate cell fractions differences in Sed and ExT mouse PV determined by CIBERSORTx. Fractions were calculated by estimating relative subsets of RNA transcripts in the RNAseq and using cell matrix from publicly available snRNAseq dataset (GSE183310) as a reference. P-values were identified with χ2-squared test. **D**, Representative 5 µM section of ExT mouse PV-LA junction in Xenium spatial *in situ* transcriptional study indicating heterogeneity of cell types detected in the PV-LA junction. Each color-labeled dot corresponds to a specific transcript captured from a particular location in the tissue section. 11 cell types were identified across 6 preparations and were assigned to a unique color (red –CM1; dark green – CM2; CM3 – dark blue; Macrophage – dark yellow; Fibroblasts – green; Adipocytes – pink; Endothelial cells – orange; Myofibroblasts – magenta; Endocardium – dark grey; Epithelial cells – purple; Smooth Muscle – blue). **E**, Uniform manifold approximation and projection (UMAP) representation of cell types identified across Sed (n = 3) and ExT (n = 3) PV preparations. **F**, UMAP in Sed samples (Top) and ExT (bottom). **G**, Waffle plots demonstrating mean cell fractions of a specific cell type (color coded) in Sed or ExT in LA and PV. **H**, Representative image of fibroblasts (green) and myofibroblasts (magenta) showing spatial distribution across PV regions in Sed and ExT mice. PV-specific CMs highlighted. **I**, Representative image of fibroblasts (green) and myofibroblasts (magenta) showing spatial distribution across LA of Sed and ExT mice. LA-specific CM population highlighted. **J**, Heatmaps depicting Z-scores from neighbourhood enrichment analysis, indicating spatial interactions between cell-type pairs in Sed (top) and ExT (bottom) conditions. Significant interactions (|Z| > 1.96, p < 0.05) show enrichment or depletion, with non-significant interactions (|Z| ≤ 1.96, p > 0.05) reflecting random spatial patterns.

### Molecular landscape of the PV-LA junction at subcellular resolution

We applied subcellular-resolution spatial transcriptomics (10x Genomics Xenium ISS platform) to 5 μm transverse sections of formalin-fixed, paraffin-embedded (FFPE) biopsies of the mouse PV-LA junction (**Figure 5D**; n=4 Sed and n=5 ExT preparations). We supplemented the 10x Genomics’ standard mouse profiling panel with a 100-gene custom panel with padlock probes for a selection of transcripts relating to ion channels, inflammation and the extracellular matrix associated with AF, resulting in 450 mRNA transcript assays (**Supplementary Table S4**) which were hybridised, amplified, and then sequenced *in situ* within each sample. Individual cells were detected by nuclear DAPI staining and cell boundaries were segmented by expanding outwards until 5 μm or the boundary of another cell was reached (see **Supplemental Methods**). This yielded 96,879 individual cells across 3 Sed and 3 ExT PV preparations that passed quality control. When mapped to histology data, the spatial transcriptomics approach clearly resolved the venous, cardiomyocyte, and fibrotic regions of the PV (**Supplemental Figure S6**). Next, we clustered cells based on their transcriptional profiles and compared cell type composition between samples after normalising transcript counts using scale factor normalisation. Examination of differentially expressed marker genes in each cluster identified cardiomyocyte populations expressing ion channels (e.g. *Kcnj3*, *Kcnj2*, Kcnd2, *Kcna4*, *Cacna1c*, *Cacna1d*, *Hcn1* and *Hcn4*), fibroblasts (marked by *Htra3*, *Pcdh9*, *Pi16*, *Col3a1*, *Col1a2, Mmp2*, *Htra3*), smooth muscle cells (marked by *Myh11*, *Acta2*, *Myl9*, *Tagln*, *Cald1*) and endothelial cells with high expression of *Rbp7*, *Tcf15*, *Gpihbp1*, *Timp4* and *Mgll*. We then used canonical cell type markers derived from published mouse PV and LA single cell RNAseq datasets (utilised in CIBERSORTx analysis described above) and label transfer-based methods to collate cells into archetypical clusters (**Figure 5E**) including three cardiomyocyte clusters, fibroblasts, macrophages, smooth muscle cells, endothelial cells etc.) that could be directly compared across specimens. Uniform Manifold Approximation and Projection (UMAP) differentiated Sed and ExT cells into 11 clusters per specimen (**Figure 5F**). We investigated the impact of exercise training on the relative proportion of major cell types in PV and LA regions by applying a χ^2^ test to specific cell type counts normalised to the total cell count per replicate (**Figure 5G**). This approach indicated distinct variability in the 3 detected cardiomyocyte cell types (considered in detail in **Figure 6** and associated text below). The relative abundance of macrophages was not modified by region or training status. Beside macrophages, other immune cells were detected inconsistently across 6 sections: neutrophils (expressing *Cxcr2*, *Csf3r*, *Cd300lf*, *Ptprc*) present in one ExT sample, T cells (expressing *Cd3d*, *Ms4a6b*, *Ctla4*, *Epsti1*) and monocytes (expressing *Ms4a6c*, *Alox5ap*, *Cybb*, *Lst1*, *Apoc2*) in one Sed sample. Consistent with enhanced ECM deposition in the PV-LA junction described in **Figure 3J**, myofibroblasts represented an increased proportion of overall cell number in ExT samples in LA but not PV regions (**Figure 5H**, PV: 0.8% in Sed *vs*. 5.2% in ExT; P=0.161; LA: 3.9% in Sed *vs*. 18.72 in ExT; p=0.001). Fibroblasts were significantly more enriched ExT PV *vs* Sed PV regions **Figure 5I**, increasing from 17.1% to 27.4% p=0.023). Given the detected changes in cell composition, we hypothesised disparities in cell type spatial organisation between ExT and Sed datasets. The single-cell resolution of our data permitted neighbourhood analysis to quantify spatial proximity between different cell types (**Figure 5J**). In both datasets, multiple interactions illustrated statistically significant enrichment or depletion, indicating deviations from random spatial associations. Notably, in ExT datasets the interaction between fibroblasts and cardiomyocytes (CM1) was enriched (FB2-CM1, Z = 19.45, p<0.001). This result implies a strong spatial correlation, likely underpinning the observed increase in ECM deposition seen in **Figure 3J**. In the Sed dataset, the interaction between fibroblasts and CM1 exhibited the highest level of depletion, with a Z-score of -72.36 (p < 0.001), indicating spatial de-coupling. Furthermore, CM1-myofibroblast interactions were also depleted in the Sed dataset with a Z-score of -16.65 (p<0.001), implying fewer interactions, a more static distribution, and an anti-fibrotic milieu in Sed compared to ExT hearts.

**Figure 6.**
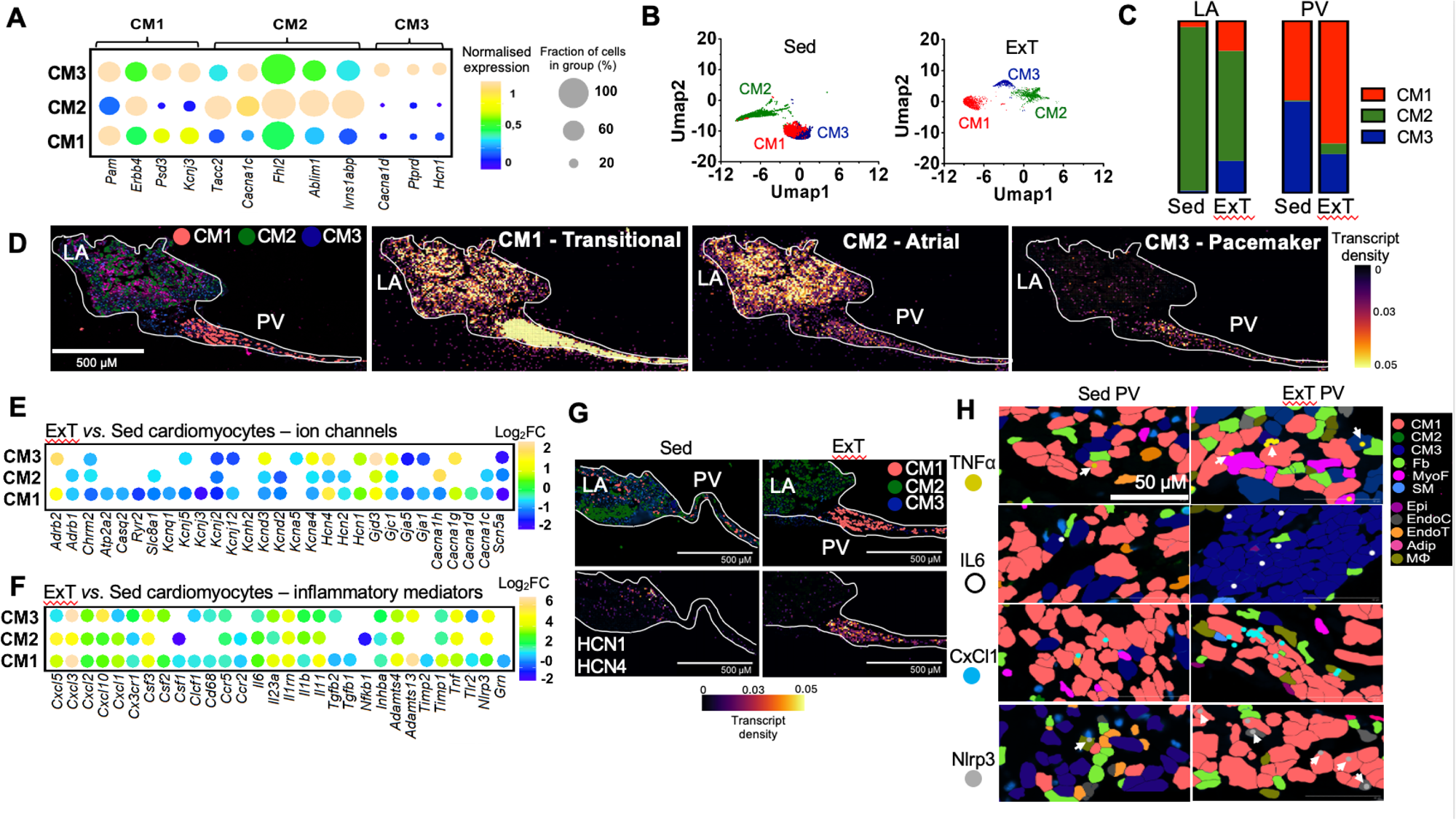
Ion channel remodeling and pro-inflammatory signaling in ExT PV cardiomyocytes. **A**, Dot plot representing stratification of CM subtypes from Xenium *in situ* experiment in n=3 Sed and n=3 ExT PV-LA preparations based on published (GSE183310) markers for mouse PV and LA. Dot sizes represent % of cells in a subtype expressing a given gene, colors indicate mean normalized expression across all 3 subtypes. **B**, UMAP representation of CMs subtypes identified by Xenium spatial transcriptomics in Sed (Left) and ExT (Right). **C**, Stacked bar charts displaying CMs subtype percentage in LA or PV of Sed or ExT mice. **D**, Xenium image of LA-PV junction (left panel) and transcript density plots for CMs markers giving density of marker genes in tissue per bin (10 × 10 μm in size). Color gradient in legend indicates the number of corresponding transcripts captured in each bin. Highest transcript density (0.05) given in yellow and lowest (0) in purple. **E-F**, Dot plot of transcripts encoding key ion channels (top) and proteins related to inflammation (bottom) showing significant (Wilcoxon test, FDR<0.05) dysregulation between ExT and Sed groups in Xenium experiments. Color legend describes up-regulated transcripts with (green-yellow-beige) and down-regulated transcripts with (green-cyan-blue). Fold changes given in logarithmic scale. **G**, Representative spatial distribution of cardiomyocyte subtypes in LA-PV junction (top) and corresponding *Hcn* channel transcript density plots (bottom). Color gradient in legend indicates the number of corresponding transcripts captured in each bin. Highest transcript density (0.05) given in yellow and lowest (0) in purple. **H**, Representative spatial distribution of selected pro-inflammatory transcripts in PV cell types of Sed and ExT mice: TNF*α* (yellow), IL-6 (white), CxCl1 (cyan) and NLRP3 (grey). Cell types are highlighted with color coding in line with previous figure and described in panel.

### Ion channel remodelling and enhanced pro-inflammatory signalling in ExT PV cardiomyocytes

Given the training-induced electrophysiological changes described in **Figures 3** and **4**, spatial heterogeneity and ion channel remodelling in cardiomyocyte populations was explored in further detail. As given above, sub-clustering marker genes selected from previous snRNAseq analysis^40^ segregated cardiomyocytes into 3 distinct clusters CM1, CM2 and CM3 (**Figure 6A)**, and the separation between the CM1 and CM3 populations was more distinct in the ExT group (**Figure 6B**). Analysis of relative ion channel expression in the 3 groups (**Supplemental Table S4**) determined that genes encoding pacemaking ion channels *Hcn1*, *Hcn4* and *Cacna1d* were highly expressed in CM3 cells, suggestive of automaticity in this population. CM2 cells exhibited an atrial-like phenotype (e.g. with higher expression of *Cacna1c* underlying Ca_v_1.2) whereas CM1 had an expression pattern transitional between CM2 and CM3 (detailed comparison between the populations is given in **Supplemental Figure S7, Supplemental Table S4** and accompanying Supplemental Discussion).

Transcripts visualized through Xenium Explorer revealed clear and consistent distribution gradients of CM subtypes in all preparations. An exemplar density map of key marker transcripts in a Sed PV preparation is given in **Figure 6D** showing enrichment of *Cacna1d*, *Ptprd* and *Hcn1* in the (pacemaker-like) CM3 population in the PV region (right panel) and *Cacna1c*, *Tacc2*, *Fhl2*, *Ablim1*, *Ivns1abp*-expressing atrial-like CM2 population predominant in the LA (central panel). Summary data pooling such information from all preparations is given in **Figure 6C** and shows that transitional-like CM1 populations are present in both LA and PV, whereas atrial-like CM2 predominate in the LA (95% in Sed LA) and pacemaker-like CM3 in the PV (53.2% in Sed LA). In agreement with our CIBERSORTx findings (**Figure 5C**), the atrial-like CM2 population was enhanced in ExT PV (0.7% in Sed PV *vs*. 6.2% in ExT PV) whereas transitional-like CM1 and pacemaker-like CM3 populations in the LA also significantly increased in ExT-LA by 13.9% (χ^2^ test, p=0.004) and 17.3% (χ^2^ test, p=1.73 x 10^-5^) respectively. These data determine a distinct spatial organisation and composition of cardiomyocytes within the PV-LA junction in the trained heart. Analysis of ion channels underpinning key ionic currents in PV and LA myocytes (**Figure 6E**) revealed multiple changes to ion channel gene transcription in ExT myocytes with the following notable differences: (i) enhanced expression of transcripts underlying pacemaking ion channels HCN1, HCN4, Ca_v_1.3 and Ca_v_3.1 in CM1 and CM3 populations in ExT samples (**Figure 6E,G, Supplemental Figure S8**), consistent with increased PV spontaneous activity and excitability (**Figure 3G**), increased frequency of pacemaker-like APs in the trained PV (**Figure 4E**) and in partial agreement with RNAseq data determining enhanced HCN4 expression (**Figure 5A);** (ii) significant downregulation of *Scn5a* and *Gja1* in all CM types consistent with observed conduction slowing (**Figure 3I**); (iii) significant downregulation of *Cacna1c* in CM1 and CM2 which could lead to shorter APDs **(Supplemental Figure S4A)** and impaired *I*_Ca,L_, a hallmark of atrial remodelling in AF. Other training-induced changes to ion channels and Ca^2+^ handling proteins are considered further in **Supplemental Figure S7, Supplemental Table S4** and accompanying Supplemental Discussion.

Surprisingly, analysis of all differentially expressed transcripts revealed a striking upregulation in many of the measured immune-inflammatory mediators *within* ExT cardiomyocytes (**Figure 6F**). The subcellular resolution enabled by Xenium allowed visualisation and comparison of transcripts related to inflammation and the fibrotic response localised to individual cardiomyocytes (**Figure 6F**,**H**) and in this study we observed that ExT PV cardiomyocytes presented with significant upregulation of genes encoding NACHT, LRR, and PYD domain containing protein 3 (*Nlrp3*) and the transcription factor NFκB (*Nfkb*) pro-inflammatory and pro-fibrotic cytokines (TNFα, Il6, Il1β, Il11), C-X-C motif chemokine ligands involved in immune cell recruitment (CXCL3, 5 and 19), chemokine receptors that recruit monocytes and macrophages to inflamed tissue, C-C chemokine receptors (CCR2 and CCR5) as well as master regulators of fibrosis (TGFβ1, TGFβ2, TIMP1 and TIMP2), all of which have known associations with AF.^19,20,45,49–53^ These findings highlight dynamic regulation of genes related to inflammation in PV cardiomyocytes and are in line with published data^19,45,46^ mechanistically linking pro-inflammatory signalling in cardiomyocytes with electrical and structural atrial remodelling as described in **Figures 3** and **4**.

## DISCUSSION

This study is the first demonstration that sustained high-intensity endurance exercise results in pro-arrhythmic remodelling of the PV and PV-LA junction, clarifying the mechanistic basis for AF susceptibility in endurance athletes, and explaining the high levels of treatment success by PV isolation in human athletes. For the first time, we have investigated a large animal model of the training-induced AF substrate, which coupled with mouse studies, identified a range of electrical and structural remodelling changes in the PV-LA junction known to increase AF propensity – these include intrinsic bradycardia, enhanced PV myocyte excitability and electrophysiological remodelling, AP heterogeneity and conduction slowing. Taken together these will predispose the athlete’s heart to initiation and anchoring of a macro-reentrant arrhythmia. We also present the first spatial analysis of the molecular landscape of the PV-LA junction that localises cardiomyocyte populations to distinct niches and dissects new roles for ion channel remodelling and pro-inflammatory signalling in PV cardiomyocytes as mediators in the heightened AF susceptibility of the athlete’s heart. The granularity of our single cell analyses reveals a greater degree of functional and transcriptomic heterogeneity within the PV-LA junction than recognized in current models, with implications for understanding of AF pathogenesis that may extend to the general population.

Since human mapping studies have identified the PVs as commonest sites of AF initiation, much effort has been devoted towards understanding the pro-arrhythmogenic characteristics of the PVs and the PV-LA junction. Unique ion-channel distribution and transmembrane conductance,^54^ spontaneous Ca^2+^ leak and oscillations,^55^ arrhythmogenic responses to local autonomic activity^56^ and tissue anisotropy^57^ are understood to precipitate aberrant PV electrical activity. However, there is a paucity of studies examining in depth the mechanisms or pathology by which this baseline but presumably latent PV arrhythmogenic activity becomes manifest in the AF patient or in translatable models of the AF substrate. In this study we identified increased automaticity, triggered activity and slowed conduction in the ExT PV, and interrogated the underlying mechanisms. Our unbiased approaches demonstrated that cardiomyocyte populations expressing a complement of pacemaking ion channels (HCN1, HCN4, Ca_v_1.3 and Ca_v_3.1) are increased in ExT PV. By facilitating automaticity in the PV myocardium and in the surrounding LA, these cardiomyocyte subtypes can mediate ectopic beat generation, constituting focal sources potentially able to initiate AF.^41^ Particularly intriguing is the chamber-specific remodelling of HCNs in response to exercise. We previously determined reduced HCN4 and *I*_f_ in the trained sinoatrial^29,31^ and atrioventricular node^32^ and contrastingly, increased ectopic HCN4 expression in the ventricles.^26^ Similarly, onset and maintenance of age-related^58^ or tachypacing-induced AF in canines^59^ have been linked to lower HCN channel expression in the sinoatrial node coupled with higher expression in the PV and LA. Further investigation of the epigenetic program that dictates tissue-specific expression of HCNs and other channels involved in pacemaking may reveal novel targets for suppressing ectopic activity and AF.

In addition to the PV myocytes being a source of ectopic firing, it is likely that specific electrophysiological characteristics of the PV-LA junction predispose to AF initiation in humans because ectopy from other regions of the atrium such as the crista terminalis do not commonly degenerate into AF, and in some people rapid sustained atrial tachycardia from the PVs can propagate to the LA without initiating AF.^60^ Our data indicate that ion channel changes in ExT PV myocytes, in addition to fibroblast mediated ECM deposition are involved in the conduction slowing and increased propensity to re-entry in the ExT PV-LA junction. Following exercise training there was a substantial downregulation of *Scn5a* and *Gja1* in all myocyte types and this is expected to slow AP conduction if translated into a downregulation of *I*_Na_ and electrical coupling.^61,62^ In all myocyte types there was a downregulation of *Kcnj2* – if this is translated into a decrease in the inward rectifier K^+^ current, *I*_K,1_, more positive diastolic potentials and a further decrease in *I*_Na_ and conduction velocity as a result of inactivation of *I*_Na_ can be anticipated. Interestingly, there was a substantial downregulation of transcripts for parasympathetic signalling i.e. the M2 muscarinic receptor and the Kir3.1 subunit of the ACh-activated K^+^ channel (*Chrm2* and *Kcnj3*) in the PV myocytes following exercise training.

Our discovery of enhanced pro-inflammatory signalling within ExT PV myocytes adds to a growing body of evidence supporting exploration of inflammation as a therapeutic target for AF.^42^ Recent studies have determined specific contributions of the pro-inflammatory cytokine TNFα,^20^ the IL-1β-maturation platform and the NLRP3 inflammasome^45^ to electrophysiological remodelling and ECM deposition in AF. Among the myriad AF-associated cytokine, chemokine, profibrotic and proinflammatory signalling molecules we have shown to be activated in ExT PV cardiomyocytes, a specific role for the mechanosensitive and pro-inflammatory cytokine TNFα in mediating athletic AF susceptibility is emerging: TNFα elevation has been reported in marathon runners,^63^ and elevated circulating TNFα levels are correlated with post exercise cardiac dysfunction in athletes competing in triathlons.^64^ Backx and colleagues demonstrated that genetic ablation or pharmacological inhibition of TNFα prevented exercise-induced adverse atrial remodelling and AF promotion in mice.^19,20^ The upstream determinants of TNFα induction in athletic hearts have yet to be determined but atrial stetch, long associated with LA and PV remodelling in AF^65^ is a possible mechanism. Myocardial stretch accompanies chamber dilation, and LA volumes are estimated to be at least 30% greater in athletes than non-athletes.^66^ Mechanical stretch resulted in enhanced PV automaticity and the occurrence of afterdepolarisations in a manner dependent on the intensity of the stretch applied. In other studies, stretch increased PV expression of TNFα^67^ and other pro-inflammatory proteins including Il-6, CxCl1 and Cx3Cl1 that were upregulated in ExT PV. Mechanistically, cyclic mechanical stretch activated NFκB in cardiomyocytes, a direct transcriptional regulator of TNFα, also enhanced in ExT PV myocytes (**Figure 6F**).^68^ Further interrogation of the precise mechanistic links between mechanical forces, TNFα activation and electrical remodelling are now required to guide the rational design of new strategies for AF prevention in the athletic population.

## Limitations

Our data cannot be directly extrapolated to AF in the human athlete in which structural and electrophysiological changes presumably develop over decades of exercise training. Nevertheless, we have investigated 4 months of exercise training (analogous to ∼5 years in human terms) in a canine model, with demonstrated anatomic, hemodynamic and electrophysiological similarities to the human heart, as a step in this direction. We did not assess PV electrical remodelling at the protein and ionic current level. Our intracellular recordings coupled with machine learning and spatial transcriptional data demonstrate that PV myocytes cannot be regarded or investigated as a singular monolithic population of cells. Thus, adequately powered studies using standard techniques (e.g. Western blotting and voltage clamp) to probe exercise-induced changes in protein levels and ionic currents in isolated cardiomyocyte subpopulations require approaches (i.e. transgenic-based subtype targeting) and consequently numbers of trained animals well beyond the present scope. The time course of remodelling changes was not investigated, and it is at present unknown whether electrical and structural remodelling proceed simultaneously, or whether they reverse with detraining. Despite these limitations, our data shed new light on the fundamental biology of PV myocytes and on PV-mediated mechanisms responsible for the increased prevalence of AF in athletes.

## REFERENCES

1. D’Souza A, Trussell T, Morris GM, Dobrzynski H, Boyett MR. Supraventricular arrhythmias in athletes: Basic mechanisms and new directions. Physiology (Bethesda). 2019;34:314–326. doi: 10.1152/physiol.00009.2019

2. Heidbuchel H, Hoogsteen J, Fagard R, Vanhees L, Ector H, Willems R, Van Lierde J. High prevalence of right ventricular involvement in endurance athletes with ventricular arrhythmias. Role of an electrophysiologic study in risk stratification. European Heart Journal. 2003;24:1473–1480.

3. Heidbuchel H, Prior DL, La Gerche A. Ventricular arrhythmias associated with long-term endurance sports: What is the evidence? British Journal of Sports Medicine. 2012;46 Suppl 1:i44–50. doi: 10.1136/bjsports-2012-091162

4. Karjalainen J, Kujala UM, Kaprio J, Sarna S, Viitasalo M. Lone atrial fibrillation in vigorously exercising middle aged men: Case-control study. British Medical Journal. 1998;316:1784–1785.

5. Mont L, Sambola A, Brugada J, Vacca M, Marrugat J, Elosua R, Pare C, Azqueta M, Sanz G. Long-lasting sport practice and lone atrial fibrillation. European Heart Journal. 2002;23:477–482. doi: 10.1053/euhj.2001.2802

6. Aizer A, Gaziano JM, Cook NR, Manson JE, Buring JE, Albert CM. Relation of vigorous exercise to risk of atrial fibrillation. American Journal of Cardiology. 2009;103:1572–1577. doi: 10.1016/j.amjcard.2009.01.374

7. Elosua R, Arquer A, Mont L, Sambola A, Molina L, Garcia-Moran E, Brugada J, Marrugat J. Sport practice and the risk of lone atrial fibrillation: A case-control study. International Journal of Cardiology. 2006;108:332–337. doi: 10.1016/j.ijcard.2005.05.020

8. Grimsmo J, Grundvold I, Maehlum S, Arnesen H. High prevalence of atrial fibrillation in long-term endurance cross-country skiers: Echocardiographic findings and possible predictors--a 28-30 years follow-up study. European journal of cardiovascular prevention and rehabilitation. 2010;17:100–105. doi: 10.1097/HJR.0b013e32833226be

9. Heidbuchel H, Anne W, Willems R, Adriaenssens B, Van de Werf F, Ector H. Endurance sports is a risk factor for atrial fibrillation after ablation for atrial flutter. International Journal of Cardiology. 2006;107:67–72. doi: 10.1016/j.ijcard.2005.02.043

10. Hoogsteen J, Schep G, Van Hemel NM, Van Der Wall EE. Paroxysmal atrial fibrillation in male endurance athletes. A 9-year follow up. Europace. 2004;6:222–228. doi: 10.1016/j.eupc.2004.01.004

11. Molina L, Mont L, Marrugat J, Berruezo A, Brugada J, Bruguera J, Rebato C, Elosua R. Long-term endurance sport practice increases the incidence of lone atrial fibrillation in men: A follow-up study. Europace. 2008;10:618–623. doi: 10.1093/europace/eun071

12. Mont L, Tamborero D, Elosua R, Molina I, Coll-Vinent B, Sitges M, Vidal B, Scalise A, Tejeira A, Berruezo A, et al. Physical activity, height, and left atrial size are independent risk factors for lone atrial fibrillation in middle-aged healthy individuals. Europace. 2008;10:15–20. doi: 10.1093/europace/eum263

13. Baldesberger S, Bauersfeld U, Candinas R, Seifert B, Zuber M, Ritter M, Jenni R, Oechslin E, Luthi P, Scharf C, et al. Sinus node disease and arrhythmias in the long-term follow-up of former professional cyclists. Eur Heart J. 2008;29:71–78.

14. Guasch E, Mont L. Diagnosis, pathophysiology, and management of exercise-induced arrhythmias. Nature Reviews Cardiology. 2017;14:88–101. doi: 10.1038/nrcardio.2016.173

15. Myrstad M, Nystad W, Graff-Iversen S, Thelle DS, Stigum H, Aaronaes M, Ranhoff AH. Effect of years of endurance exercise on risk of atrial fibrillation and atrial flutter. American Journal of Cardiology. 2014;114:1229–1233. doi: 10.1016/j.amjcard.2014.07.047

16. Newman W, Parry-Williams G, Wiles J, Edwards J, Hulbert S, Kipourou K, Papadakis M, Sharma R, O’Driscoll J. Risk of atrial fibrillation in athletes: A systematic review and meta-analysis. British Journal of Sports Medicine. 2021;55:1233–1238. doi: 10.1136/bjsports-2021-103994

17. Benito B, Gay-Jordi G, Serrano-Mollar A, Guasch E, Shi Y, Tardif JC, Brugada J, Nattel S, Mont L. Cardiac arrhythmogenic remodeling in a rat model of long-term intensive exercise training. Circulation. 2011;123:13–22. doi: 10.1161/CIRCULATIONAHA.110.938282

18. Guasch E, Benito B, Qi X, Cifelli C, Naud P, Shi Y, Mighiu A, Tardif JC, Tadevosyan A, Chen Y, et al. Atrial fibrillation promotion by endurance exercise: Demonstration and mechanistic exploration in an animal model. Journal of the American College of Cardiology 2013;62:68–77. doi: 10.1016/j.jacc.2013.01.091

19. Aschar-Sobbi R, Izaddoustdar F, Korogyi AS, Wang Q, Farman GP, Yang F, Yang W, Dorian D, Simpson JA, Tuomi JM, et al. Increased atrial arrhythmia susceptibility induced by intense endurance exercise in mice requires TNFalpha. Nature Communications. 2015;6:6018. doi: 10.1038/ncomms7018

20. Lakin R, Polidovitch N, Yang S, Guzman C, Gao X, Wauchop M, Burns J, Izaddoustdar F, Backx PH. Inhibition of soluble tnfalpha prevents adverse atrial remodeling and atrial arrhythmia susceptibility induced in mice by endurance exercise. Journal of Molecular and Celularl Cardiology. 2019;129:165–173. doi: 10.1016/j.yjmcc.2019.01.012

21. Haissaguerre M, Jais P, Shah DC, Takahashi A, Hocini M, Quiniou G, Garrigue S, Le Mouroux A, Le Metayer P, Clementy J. Spontaneous initiation of atrial fibrillation by ectopic beats originating in the pulmonary veins. The New England Journal of Medicine. 1998;339:659–666. doi: 10.1056/NEJM199809033391003

22. Kumagai K, Ogawa M, Noguchi H, Yasuda T, Nakashima H, Saku K. Electrophysiologic properties of pulmonary veins assessed using a multielectrode basket catheter. Journal of the American College of Cardiology. 2004;43:2281–2289. doi: 10.1016/j.jacc.2004.01.051

23. Hocini M, Ho SY, Kawara T, Linnenbank AC, Potse M, Shah D, Jais P, Janse MJ, Haissaguerre M, De Bakker JM. Electrical conduction in canine pulmonary veins: Electrophysiological and anatomic correlation. Circulation. 2002;105:2442–2448. doi: 10.1161/01.cir.0000016062.80020.11

24. Lee G, Spence S, Teh A, Goldblatt J, Larobina M, Atkinson V, Brown R, Morton JB, Sanders P, Kistler PM, et al. High-density epicardial mapping of the pulmonary vein-left atrial junction in humans: Insights into mechanisms of pulmonary vein arrhythmogenesis. Heart Rhythm. 2012;9:258–264. doi: 10.1016/j.hrthm.2011.09.010

25. Mandsager KT, Phelan DM, Diab M, Baranowski B, Saliba WI, Tarakji KG, Jaber WA, Kanj M, Tchou P, Lindsay BD, et al. Outcomes of pulmonary vein isolation in athletes. Journal of the American College of Cardiology:Clinical Electrophysiology 2020;6:1265–1274. doi: 10.1016/j.jacep.2020.05.009

26. Polyak A, Topal L, Zombori-Toth N, Toth N, Prorok J, Kohajda Z, Deri S, Demeter-Haludka V, Hegyi P, Venglovecz V, et al. Cardiac electrophysiological remodeling associated with enhanced arrhythmia susceptibility in a canine model of elite exercise. Elife. 2023;12. doi: 10.7554/eLife.80710

27. Wright KN, Hines DA, Bright JM. Cardiac electrophysiologic measurements in dogs before and after intravenous administration of atropine and propranolol. American Journal of Veterinary Research. 1996;57:1695–1701.

28. Elvan A, Wylie K, Zipes DP. Pacing-induced chronic atrial fibrillation impairs sinus node function in dogs. Electrophysiological remodeling. Circulation. 1996;94:2953–2960. doi: 10.1161/01.cir.94.11.2953

29. D’Souza A, Pearman CM, Wang Y, Nakao S, Logantha S, Cox C, Bennett H, Zhang Y, Johnsen AB, Linscheid N, et al. Targeting mir-423-5p reverses exercise training-induced hcn4 channel remodeling and sinus bradycardia. Circulation Research. 2017;121:1058–1068. doi: 10.1161/CIRCRESAHA.117.311607

30. Bidaud I, D’Souza A, Forte G, Torre E, Greuet D, Thirard S, Anderson C, Chung You Chong A, Torrente AG, Roussel J, et al. Genetic ablation of g protein-gated inwardly rectifying k(+) channels prevents training-induced sinus bradycardia. Frontiers in Physiology. 2020;11:519382. doi: 10.3389/fphys.2020.519382

31. D’Souza A, Bucchi A, Johnsen AB, Logantha SJ, Monfredi O, Yanni J, Prehar S, Hart G, Cartwright E, Wisloff U, et al. Exercise training reduces resting heart rate via downregulation of the funny channel hcn4. Nature Communications. 2014;5:3775. doi: 10.1038/ncomms4775

32. Mesirca P, Nakao S, Nissen SD, Forte G, Anderson C, Trussell T, Li J, Cox C, Zi M, Logantha SJR, et al. Intrinsic electrical remodeling underlies atrioventricular block in athletes. Circulation Research. 2021;129:e1–e20. doi: 10.1161/CIRCRESAHA.119.316386

33. Kemi OJ, Loennechen JP, Wisloff U, Ellingsen O. Intensity-controlled treadmill running in mice: Cardiac and skeletal muscle hypertrophy. Journal of Applied Physiology (1985). 2002;93:1301–1309. doi: 10.1152/japplphysiol.00231.2002

34. Tikhomirov R, Oakley R, Anderson C, Yaar S, Mesirca P, Torre E, Al Othman S, Smith M, Donaldson I, Soattin L, et al. Cardiac GR determines the diurnal rhythm in ventricular arrhythmia susceptibility. Circulation Research. 2024. doi: 10.1161/CIRCRESAHA.123.323464

35. Li D, Fareh S, Leung TK, Nattel S. Promotion of atrial fibrillation by heart failure in dogs: Atrial remodeling of a different sort. Circulation. 1999;100:87–95. doi: 10.1161/01.cir.100.1.87

36. Tsuneoka Y, Kobayashi Y, Honda Y, Namekata I, Tanaka H. Electrical activity of the mouse pulmonary vein myocardium. Journal of Pharmacological Sciences. 2012;119:287–292. doi: 10.1254/jphs.12062sc

37. Miyauchi Y, Hayashi H, Miyauchi M, Okuyama Y, Mandel WJ, Chen PS, Karagueuzian HS. Heterogeneous pulmonary vein myocardial cell repolarization implications for reentry and triggered activity. Heart Rhythm. 2005;2:1339–1345. doi: 10.1016/j.hrthm.2005.09.015

38. Lagomarsino-Oneto D, De Leo A, Stocchino A, Cucco A. Unraveling the non-homogeneous dispersion processes in ocean and coastal circulations using a clustering approach. Geophysical Research Letters. 2024;51:e2023GL107900. doi: 10.1029/2023GL107900

39. Bishop CM. Pattern recognition and machine learning. Springer New York; 2016.

40. Steimle JD, Grisanti Canozo FJ, Park M, Kadow ZA, Samee MAH, Martin JF. Decoding the pitx2-controlled genetic network in atrial fibrillation. Journal of Clinical Investigation Insight. 2022;7. doi: 10.1172/jci.insight.158895

41. Chen YJ, Chen SA, Chang MS, Lin CI. Arrhythmogenic activity of cardiac muscle in pulmonary veins of the dog: Implication for the genesis of atrial fibrillation. Cardiovascular Research. 2000;48:265–273. doi: 10.1016/s0008-6363(00)00179-6

42. Dobrev D, Heijman J, Hiram R, Li N, Nattel S. Inflammatory signalling in atrial cardiomyocytes: A novel unifying principle in atrial fibrillation pathophysiology. Nature Reviews Cardiology. 2023;20:145–167. doi: 10.1038/s41569-022-00759-w

43. Flannery MD, Kalman JM, Sanders P, La Gerche A. State of the art review: Atrial fibrillation in athletes. Heart Lung Circulation. 2017;26:983–989. doi: 10.1016/j.hlc.2017.05.132

44. Sellami M, Al-Muraikhy S, Al-Jaber H, Al-Amri H, Al-Mansoori L, Mazloum NA, Donati F, Botre F, Elrayess MA. Age and sport intensity-dependent changes in cytokines and telomere length in elite athletes. Antioxidants (Basel). 2021;10. doi: 10.3390/antiox10071035

45. Yao C, Veleva T, Scott L, Jr., Cao S, Li L, Chen G, Jeyabal P, Pan X, Alsina KM, Abu-Taha ID, et al. Enhanced cardiomyocyte nlrp3 inflammasome signaling promotes atrial fibrillation. Circulation. 2018;138:2227–2242. doi: 10.1161/CIRCULATIONAHA.118.035202

46. Zhang YL, Cao HJ, Han X, Teng F, Chen C, Yang J, Yan X, Li PB, Liu Y, Xia YL, et al. Chemokine receptor cxcr-2 initiates atrial fibrillation by triggering monocyte mobilization in mice. Hypertension. 2020;76:381–392. doi: 10.1161/HYPERTENSIONAHA.120.14698

47. Steimle JD, Grisanti Canozo FJ, Park M, Kadow ZA, Samee MAH, Martin JF. Decoding the pitx2-controlled genetic network in atrial fibrillation. Journal of Clinical Investigation: Insight. 2022;7. doi: 10.1172/jci.insight.158895

48. Newman AM, Steen CB, Liu CL, Gentles AJ, Chaudhuri AA, Scherer F, Khodadoust MS, Esfahani MS, Luca BA, Steiner D, et al. Determining cell type abundance and expression from bulk tissues with digital cytometry. Nature Biotechnology. 2019;37:773–782. doi: 10.1038/s41587-019-0114-2

49. Gao G, Dudley SC, Jr. Redox regulation, nf-kappab, and atrial fibrillation. Antioxidants and Redox Signalling. 2009;11:2265–2277. doi: 10.1089/ars.2009.2595

50. Zhou X, Liu H, Feng F, Kang GJ, Liu M, Guo Y, Dudley SC, Jr. Macrophage il-1beta mediates atrial fibrillation risk in diabetic mice. Journal of Clinical Investigation: Insight. 2024;9. doi: 10.1172/jci.insight.171102

51. Huang M, Huiskes FG, de Groot NMS, Brundel B. The role of immune cells driving electropathology and atrial fibrillation. Cells. 2024;13. doi: 10.3390/cells13040311

52. Lai YJ, Tsai FC, Chang GJ, Chang SH, Huang CC, Chen WJ, Yeh YH. Mir-181b targets semaphorin 3a to mediate tgf-beta-induced endothelial-mesenchymal transition related to atrial fibrillation. Journal of Clinical Investigation. 2022;132. doi: 10.1172/JCI142548

53. Liu Y, Xu B, Wu N, Xiang Y, Wu L, Zhang M, Wang J, Chen X, Li Y, Zhong L. Association of mmps and timps with the occurrence of atrial fibrillation: A systematic review and meta-analysis. Canadian Journal of Cardiology. 2016;32:803–813. doi: 10.1016/j.cjca.2015.08.001

54. Ehrlich JR, Cha TJ, Zhang L, Chartier D, Melnyk P, Hohnloser SH, Nattel S. Cellular electrophysiology of canine pulmonary vein cardiomyocytes: Action potential and ionic current properties. Journal of Physiology. 2003;551:801–813. doi: 10.1113/jphysiol.2003.046417

55. Chou CC, Nihei M, Zhou S, Tan A, Kawase A, Macias ES, Fishbein MC, Lin SF, Chen PS. Intracellular calcium dynamics and anisotropic reentry in isolated canine pulmonary veins and left atrium. Circulation. 2005;111:2889–2897. doi: 10.1161/CIRCULATIONAHA.104.498758

56. Chen PS, Chen LS, Fishbein MC, Lin SF, Nattel S. Role of the autonomic nervous system in atrial fibrillation: Pathophysiology and therapy. Circulation Research. 2014;114:1500–1515. doi: 10.1161/CIRCRESAHA.114.303772

57. Arora R, Verheule S, Scott L, Navarrete A, Katari V, Wilson E, Vaz D, Olgin JE. Arrhythmogenic substrate of the pulmonary veins assessed by high-resolution optical mapping. Circulation. 2003;107:1816–1821. doi: 10.1161/01.CIR.0000058461.86339.7E

58. Li YD, Hong YF, Zhang Y, Zhou XH, Ji YT, Li HL, Hu GJ, Li JX, Sun L, Zhang JH, et al. Association between reversal in the expression of hyperpolarization-activated cyclic nucleotide-gated (hcn) channel and age-related atrial fibrillation. Medical Science Monitoring. 2014;20:2292–2297. doi: 10.12659/msm.892505

59. Li JY, Wang HJ, Xu B, Wang XP, Fu YC, Chen MY, Zhang DX, Liu Y, Xue Q, Li Y. Hyperpolarization activated cation current (i(f)) in cardiac myocytes from pulmonary vein sleeves in the canine with atrial fibrillation. Journal of Geriatric Cardiology. 2012;9:366–374. doi: 10.3724/SP.J.1263.2012.04161

60. Morris GM, Segan L, Wong G, Wynn G, Watts T, Heck P, Walters TE, Nisbet A, Sparks P, Morton JB, et al. Atrial tachycardia arising from the crista terminalis, detailed electrophysiological features and long-term ablation outcomes. Journal of the American College of Cardiology : Clinical Electrophysiology. 2019;5:448–458. doi: 10.1016/j.jacep.2019.01.014

61. van Veen TA, Stein M, Royer A, Le Quang K, Charpentier F, Colledge WH, Huang CL, Wilders R, Grace AA, Escande D, et al. Impaired impulse propagation in scn5a-knockout mice: Combined contribution of excitability, connexin expression, and tissue architecture in relation to aging. Circulation. 2005;112:1927–1935. doi: 10.1161/CIRCULATIONAHA.105.539072

62. Eloff BC, Gilat E, Wan X, Rosenbaum DS. Pharmacological modulation of cardiac gap junctions to enhance cardiac conduction: Evidence supporting a novel target for antiarrhythmic therapy. Circulation. 2003;108:3157–3163. doi: 10.1161/01.CIR.0000101926.43759.10

63. Bernecker C, Scherr J, Schinner S, Braun S, Scherbaum WA, Halle M. Evidence for an exercise induced increase of tnf-alpha and il-6 in marathon runners. Scandinavian journal of medicine & science in sports. 2013;23:207–214. doi: 10.1111/j.1600-0838.2011.01372.x

64. La Gerche A, Inder WJ, Roberts TJ, Brosnan MJ, Heidbuchel H, Prior DL. Relationship between inflammatory cytokines and indices of cardiac dysfunction following intense endurance exercise. PLoS One. 2015;10:e0130031. doi: 10.1371/journal.pone.0130031

65. Gottlieb LA, Coronel R, Dekker LRC. Reduction in atrial and pulmonary vein stretch as a therapeutic target for prevention of atrial fibrillation. Heart Rhythm. 2023;20:291–298. doi: 10.1016/j.hrthm.2022.10.009

66. Iskandar A, Mujtaba MT, Thompson PD. Left atrium size in elite athletes. JACC Cardiovascular Imaging. 2015;8:753–762. doi: 10.1016/j.jcmg.2014.12.032

67. Zhao J, Nishimura Y, Kimura A, Ozawa K, Kondo T, Tanaka T, Yoshizumi M. Chemokines protect vascular smooth muscle cells from cell death induced by cyclic mechanical stretch. Scientific Reports. 2017;7:16128. doi: 10.1038/s41598-017-15867-8

68. Leychenko A, Konorev E, Jijiwa M, Matter ML. Stretch-induced hypertrophy activates nfkb-mediated vegf secretion in adult cardiomyocytes. PLoS One. 2011;6:e29055. doi: 10.1371/journal.pone.0029055

